# GWASBrewer: An R Package for Simulating Realistic GWAS Summary Statistics

**DOI:** 10.1101/2024.04.16.589571

**Authors:** Jean Morrison

## Abstract

Many statistical genetics analysis methods make use of GWAS summary statistics. Best statistical practice requires evaluating these methods in simulations against a known truth. Ideally, these simulations should be as realistic as possible. However, simulating summary statistics by first simulating individual genotype and phenotype data is extremely computationally demanding, especially when large sample sizes or many traits are required. We present `GWASBrewer`, an open source R package for direct simulation of GWAS summary statistics. We show that statistics simulated by `GWASBrewer` have the same distribution as statistics generated from individual level data, and can be produced at a fraction of the computational expense. Additionally, `GWASBrewer` can simulate standard error estimates, something that is typically not done when sampling summary statistics directly. `GWASBrewer` is highly flexible, allowing the user to simulate data for multiple traits connected by causal effects and with complex distributions of effect sizes. We demonstrate example uses of `GWASBrewer` for evaluating Mendelian randomization, polygenic risk score, and heritability estimation methods.

## 2 Introduction

In recent years there has been a proliferation of statistical methods that use effect estimates and standard errors (summary statistics) from genome-wide association studies (GWAS) to infer interesting biological parameters. These include methods for estimating heritability [1, 2, 3, 4], genetic correlation [5, 6, 7], causal effects via Mendelian randomization (MR) [8, 9, 10], and polygenic risk scores [11, 12, 13, 14]. A common challenge in the development of all methods applied to genetic data is conducting simulation studies that adequately mimic properties of real data. Overly simplistic simulations may mis-represent the expected behavior of a method. One of the most realistic strategies to generate summary statistics is the full-data simulation procedure. In this procedure, genotype data are generated by re-sampling haplotypes from a reference panel, from a population-genetic model, or from a correlated binomial distribution [15, 16, 17]. Phenotype data are generated according to particular (usually additive) genetic model and finally association estimates are computed for each variant. The resulting summary statistics can then be passed to the method or methods being evaluated.

A major drawback of the full-data simulation method is that it is computationally intensive and involves production and storage of large quantities of data. Some effort can be saved by using the same set of genotype data in every simulation replicate and only regenerating phenotype data. However, the step of computing association estimates requires performing up to millions of linear regressions. This problem is magnified when data from multiple GWAS are required. This computational demand may motivate investigators to limit the scope of their simulations or make simplifying assumptions that could alter the conclusions of the experiment. For example, many MR methods have been evaluated by generating data for a small number of pre-selected independent variants [10, 18, 19]. However, when these methods are applied in practice, variants must be selected from genome-wide data. This step can add bias to the MR estimate that is not captured by overly simplistic simulations. Another common simplification is to assume that there is no LD between variants. However, this results in much sparser genetic signal than would be expected in real data and can lead to over-optimism about the accuracy of some methods.

When evaluating methods that only require GWAS summary statistics, it is possible to directly simulate summary statistics without generating individual level data by sampling estimates from their asymptotic multivariate normal distribution. This strategy has been used by Morrison et al. [20] among others. The direct summary statistic simulation method can preserve many of the important features of real GWAS data, such as LD, with a fraction of the computational resources required by the full-data simulation method. However, despite the large number of methods developed each year for analysis of GWAS summary statistics and the common need to perform simulation and benchmarking studies, there are no well-documented R packages implementing the direct summary statistic simulation method for a range of scenarios. To fill this gap, we developed the R package `GWASBrewer`. The goal of `GWASBrewer` is to generate data that are as realistic as possible from a flexible model that can accommodate many simulation needs. Features of `GWASBrewer` include the ability to simulate data for multiple traits with specified causal relationships, for variants in linkage disequilibrium, allowing for arbitrary amounts of overlap between GWAS samples, and allowing for flexible specification of variant effect size distributions and heritability models.

In Section 3, we describe how `GWASBrewer` samples GWAS summary statistics. Section 3.3, describes the interface and specific features of `GWASBrewer`. In Section 5 we demonstrate that `GWASBrewer` accurately mimics the distribution of summary statistics obtained through full-data simulation. Finally, in Section 6, we demonstrate the use of `GWASBrewer` to test MR methods, heritability estimation methods, and polygenic risk score estimation methods.

## 3 Simulating GWAS Summary Statistics

### 3.1 Summary Statistics for a Single Trait

We first describe simulation of summary statistics for a single continuous trait, *Y*. Let *G*_1_, …, *G*_*J*_ be genetic variants with correlation *R* (frequently referred to as the linkage disequilibrium (LD) matrix). We assume that all genetic variants are bi-allelic and in Hardy-Weinberg equilibrium. Together, these assumptions mean that *G*_*j*_ is a binomial random variable with frequency *f*_*j*_, and *V ar*(*G*_*j*_) *≡ v*_*j*_ = 2*f*_*j*_(1 *− f*_*j*_).

In a GWAS, *Y* and **G** = (*G*_1_, …, *G*_*J*_) are measured for *N* individuals yielding observations (*y*_1_, **g**_1_), …, (*y*_*N*_, **g**_*N*_). The association between *G*_*j*_ and *Y* is then estimated using linear regression if *Y* is continuous, or logistic regression if *Y* is binary. Most modern GWAS include age, sex, and principal components of the genoytpe matrix as covariates, and may also use a mixed model to account for relatedness between samples. We make a simplifying assumption that GWAS samples are unrelated and sampled from a homogeneous randomly mating population and that effect estimates are estimated via simple linear regression. That is,

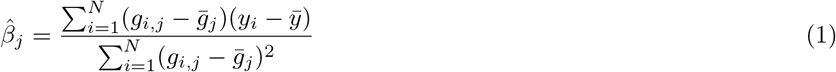

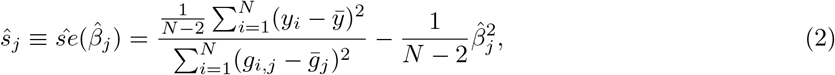

where 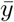 and 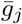 are sample means of *Y* and *G*_*j*_ respectively.

Zhu and Stephens [21] demonstrate that the joint distribution of 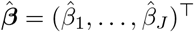 is well approximated by a multivariate normal distribution,

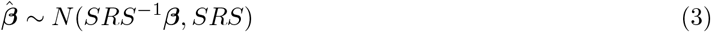

where *S* is a *J × J* diagonal matrix with *j*th diagonal entry equal to 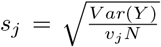 and *β* is the vector of population joint associations between variants and *Y*. For simplicity and without loss of generality, we assume that *V ar*(*Y*) = 1, so *s*_*j*_ depends only on *f*_*j*_ and *N*. The joint association between *G*_*j*_ and *Y* is the linear association conditional on all other variants. If we assume a purely additive genetic model,

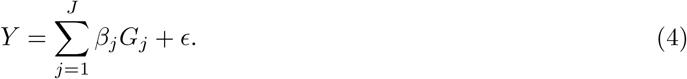

then *β* is the vector of causal effects. Joint effects are simulated from a scale-family distribution. By default, this is a point-normal distribution, however it is possible to specify other alternatives. `GWASBrewer` chooses the scale parameter so that *Y* has a user-specified expected heritability. Optionally, effects can be scaled so that heritability is exact using the `h2 exact` option. More detail on the distribution of effect sizes is given in Section 3.3.

`GWASBrewer` simulates 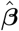 by directly sampling from the model in (3) given *β, S*, and *R*. This process is very fast when *R* is block diagonal with moderately sized blocks. Standard error estimates, *ŝ*_*j*_, can optionally be simulated as well (see Supplementary Note). The LD matrix, *R*, and allele frequencies *f*_*j*_ can be specified by the user, or `GWASBrewer` will default to generating data with no LD. The package includes one LD matrix and set of allele frequencies as built-in data objects. These were estimated from chromosome 19 in the European subset of 1000 Genomes data and broken into blocks using LDetect regions [22]. `GWASBrewer` does not require that the provided LD matrix contain the number of variants to be simulated. Instead, it repeats the provided LD pattern as many (possibly fractional) times as necessary to produce the desired number of variants. This means that genome-sized data can be generated from a smaller LD pattern and that the size of the simulated genome can be changed without generating a new reference LD object.

### 3.2 Summary Statistics for Multiple Traits

Many methods including Mendelian randomization, GenomicSEM, and cross-trait genetic correlation estimation, involve estimation of relationships between multiple traits. `GWASBrewer` can facilitate evaluation of these methods by simulating summary statistics for a a set of *K* traits, *Y*_1_, …, *Y*_*K*_ with specified genetic and environmental relationships. `GWASBrewer` can simulate summary statistics for an arbitrary number of continuous traits that are related to each other by a specified linear structural equation model (SEM) corresponding to a directed, acyclyc graph (DAG). The SEM for *K* traits is specified by the user as *D*^(*dir*)^, a *K× K* matrix of direct effects in which 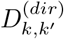 gives the direct effect of *Y*_*k*_ on *Y*_*k*_^*′*^. For example, the DAG in Figure 1 corresponds to the direct effect matrix,

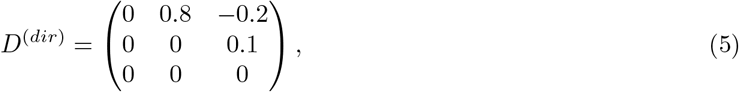

and the linear SEM,

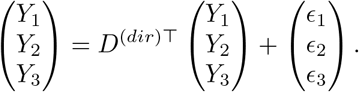

**Figure 1.**
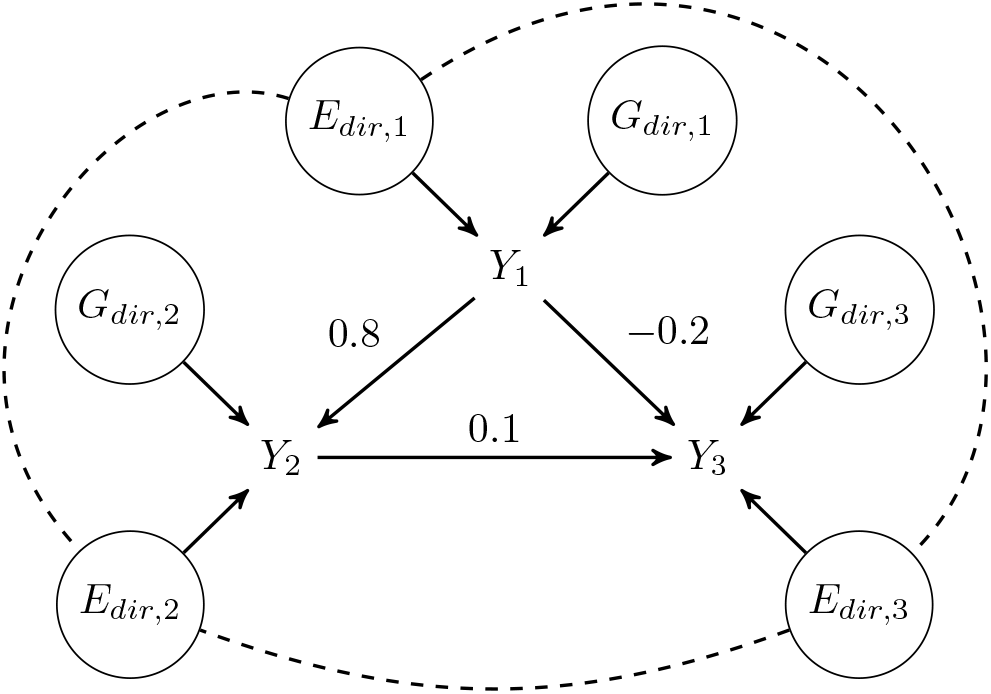
DAG Specified by the direct effects matrix in Equation (5). Nodes in circles represent direct genetic and environmental effects. Direct environmental components of traits may be correlated (dashed lines) while direct genetic components are always independent.

In this SEM, *ϵ*_*k*_, *k* = 1, 2, 3 include both direct genetic effects and direct environmental effects. In other words *ϵ*_*k*_ = *G*_*dir,k*_ + *E*_*dir,k*_ as illustrated in Figure 1. We assume that direct genetic effects are mutually independent and independent of environmental effects. However, we do not require that direct environmental effects are mutually independent of each other.

In order to simulate summary statistics for all *K* traits, `GWASBrewer` performs four steps. 1) Calculate the total variance of the direct genetic components. 2) Simulate direct effects for each variant-trait pair by sampling from up to *K* different scale-family distributions. 3) Calculate the total effect of each variant on each trait. 4) Simulate effect estimates and estimated standard errors using a multivariate extension of (3). Here we briefly describe each of these steps.

For the first and third steps, we must first compute the matrix of total trait effects. For example, in the DAG in Figure 1, the total effect of *Y*_1_ on *Y*_3_ is 0.8 *·* 0.1 *−* 0.2 =*−* 0.12. In a linear SEM corresponding to a DAG, total effects can be calculated from the matrix of direct effects as

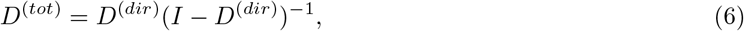

where *I* is the *K × K* identity matrix [23]. At this stage, we can also verify that the user-supplied *D*^(*dir*)^ corresponds to an acyclic graph. If this is not the case, either (*I−D*^(*dir*)^) will not be invertable or the resulting *D*^(*tot*)^ will not have 0’s on the diagonal.

Let h^2^ be a user-supplied vector of trait heritabilities. The total variance of each trait can be decomposed as *V ar*(*Y*_*k*_) = *V*_*Gdir,k*_ + *V*_*Gind,k*_ + *V*_*E,k*_, where *V*_*Gdir,k*_ is the variance due to direct genetic effects, *V*_*Gind,k*_ is the variance due to genetic effects mediated by other traits (indirect effects), and *V*_*E,k*_ is the variance due to both direct and indirect environmental effects. We assume that *V ar*(*Y*_*k*_) = 1 for all traits, so 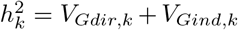. We can calculate that 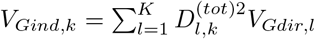. Therefore, in Step 1, we obtain the direct heritability of each trait as

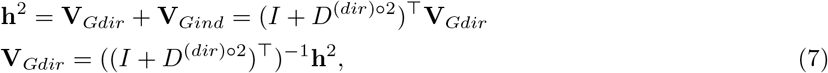

where superscript ∘2 indicates the element-wise square.

In Step 2, we sample direct effects so that the total expected variance of direct genetic effects is equal to V_*Gdir*_. Details of simulation of direct effects are given in Section 3.3. In Step 3, we use the total effects matrix to obtain the total genetic effect of each variant on each trait. Let *γ*_*j*_ be the *K*-vector giving the direct effects of variant *j* on *Y*_1_, … *Y*_*K*_. The the vector of total effects of variant *j, β*_*j*_ is

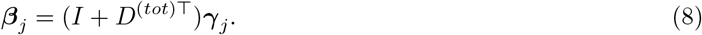

In Step 4, we simulate summary statistics using a multivariate extension of (3). The expression in (3) holds for each trait individually. However, it does not provide the covariance between effect estimates for different traits which depends on the overlap between GWAS samples and the LD between variants. Let *N*_*k*_ and *N*_*k*_^*′*^ be GWAS sample size of traits *k* and *k*^*′*^, and let *N*_*k,k*_^*′*^ be the number of overlapping samples. The correlation between 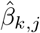 and 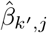 is approximately equal to 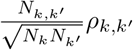, where *ρ*_*k,k*_^*′*^ is the population correlation between *Y*_*k*_ and *Y*_*k*_^*′*^ [24, 5]. More generally, the covariance between effect estimates for any pair of variants and traits is

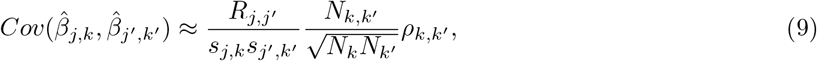

where *R*_*j,j*_^*′*^ is the correlation between variant *j* and variant *j*^*′*^ [24]. This expression allows us to sample the full *J × K* matrix of effect estimates. Estimates of the standard errors, 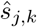 can be sampled using a multivariate generalization of the strategy used for a single trait (details in Supplementary Note). By default, the population correlation *ρ*_*k,k*_^*′*^ is calculated assuming that all direct environmental components are independent. However, the user can specify correlation between environmental components either by specifying the overall trait correlation or by specifying the overall correlation of trait environmental components.

### 3.3 Distribution of Direct Variant-Trait Effects

`GWASBrewer` supports a wide variety of genetic architectures. By default, direct effects are sparse with a proportion, *π*_*k*_, being non-zero for each trait. If *γ*_*j,k*_ is selected to be non-zero, then the standardized effect size, 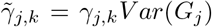, is drawn from a 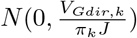 distribution, giving an expected total direct genetic variance of approximately *V*_*Gdir,k*_. This approximation assumes that most causal variants will be approximately independent, which will be true for genome-wide simulations. Using this strategy, the realized heritability will not exactly equal the input expected heritability, however, if the `h2 exact` option is used, effects will be scaled after sampling to give exactly the desired heritability.

`GWASBrewer` has options that allow the user to control both which variants are causal and the distribution of effect sizes, including permitting exact specification of causal variants and effect sizes. To control which variants are causal, the user can specify a *J× K* matrix with elements giving the probability that each variant has a direct effect on each trait. If these probabilities are set to 0 or 1, the user can deterministically select effect and non-effect variants. Alternatively, variant-specific probabilities can be used to allow the probability of being an effect variant to depend on variant-level features or to create regions of the genome with different effect densities.

To specify the effect distribution, the user can specify any function for sampling from a scale family distribution (technical details in Section 4). We provide built-in utilities for generating effect size distribution functions to sample from a mixture of normal distributions and for using a set of effects that is fixed up to a scalar. `GWASBrewer` allows for each trait to have a different effect size distribution function. It also allows the effect size distribution function to depend on variant-level annotations which can be passed to the simulation function separately.

## 4 Implementation and Interface

### 4.1 The sim_mv Function

The primary user-facing function in `GWASBrewer` is `sim_mv`. The arguments of `sim_mv` can be divided into three groups. The first describes the traits, the second describes the GWAS study design, and the third describes the distribution of direct effects of variants on traits. Trait related arguments include the matrix of direct effects, `G`, heritability, `h2`, and optionally either the observational or environmental correlation of traits, `R_obs` or `R_E`. Study design related arguments include sample size, `N`, number of variants to simulate, `J`, an optional LD pattern `R_LD` and an optional vector of allele frequencies `af`. The sample size argument can accept multiple formats and allows specification of sample overlap. Finally, effect distribution arguments include the proportion of causal variants, `pi`, an optional user-supplied effect size function or list of functions, `snp_effect_function`, and an optional dataframe of annotations that may be used by the user-specified effect function, `snp_info`. `sim_mv` produces an object of class `sim_mv` containing summary statistics as well as true marginal and joint effects, true heritability, environmental covariance, and trait correlation. A complete description of inputs and outputs for `sim_mv` is given in the Suppplementary Note.

### 4.2 Auxiliary Functions

The `GWASBrewer` package contains several functions to assist with common tasks. These include utilities for LD-pruning (`sim_ld_prune`), extracting LD proxies (`sim_ld_proxy`), and extracting the LD matrix for a set of variants (`sim_extract_ld`). `GWASBrewer` also includes two functions for generating new data with the same causal variants as an initial simulation.

It also includes two functions that make it possible to re-sample summary satistics or individual level data with the same joint effects as an initial object produced by `sim_mv`, `resample_sumstats` and `resample_inddata`. The `resample_sumstats` function could be used to simulate two separate GWAS of the same trait, possibly with different sample sizes or different LD patterns. The `resample_inddata` function generates individual level genotype and phenotype data from the effects in a previously generated simulation object. In the examples below, we show how this function can be used to test the predictive ability of a polygenic risk score. The `resample_inddata` function uses utilities from the HapSim R package [16] to simulate genotypes with the specified LD structure. It then simulates phenotypes from (4) assuming environmental effects are normally distributed with covariance given by `Sigma_E` in the original simulation object. This function requires that sample size be specified in dataframe format if there is sample overlap (i.e. matrix format is not accepted). Optionally, `resample_inddata` can efficiently calculate the linear regression summary statistics for the generated data.

## 5 Accuracy

We used simulations to verify that summary statistics generated by `GWASBrewer` follow the same distribution as summary statistics generated from individual level data. We first generated effect sizes for for 10 variants in linkage disequilibrium and two traits. The first trait has two causal variants, the second trait has three causal variants, with no common causal variants and no causal effect between traits. For both traits, causal effects from variants included explain 0.4% of trait variance. We then used `resample_sumstats` and `resample_inddata` to generate 1000 realizations of directly sampled summary statistics and 1000 realizations of summary statistics computed from individual-level data. In both cases, we specified an observational correlation between traits of 0.7. The sample size for Trait 1 was 8,000 and the sample size for Trait 2 was 10,000, with 2,000 individuals in both samples. Code for performing these simulations is provided in a supplemental file.

Figure 2 compares the mean, standard deviation, and correlation of effect estimates either directly sampled or derived from individual-level data. We find that all values are very similar between the two methods. We additionally constructed quantile-quantile (Q-Q) plots comparing the distribution of effect estimates and standard error estimates between the two methods for all variants and both traits as well as Q-Q plots comparing products of effect estimates and standard error estimates across traits. All Q-Q plots were consistent with the hypothesis that the two methods generate data with the same distribution (see Supplementary Note).

**Figure 2.**
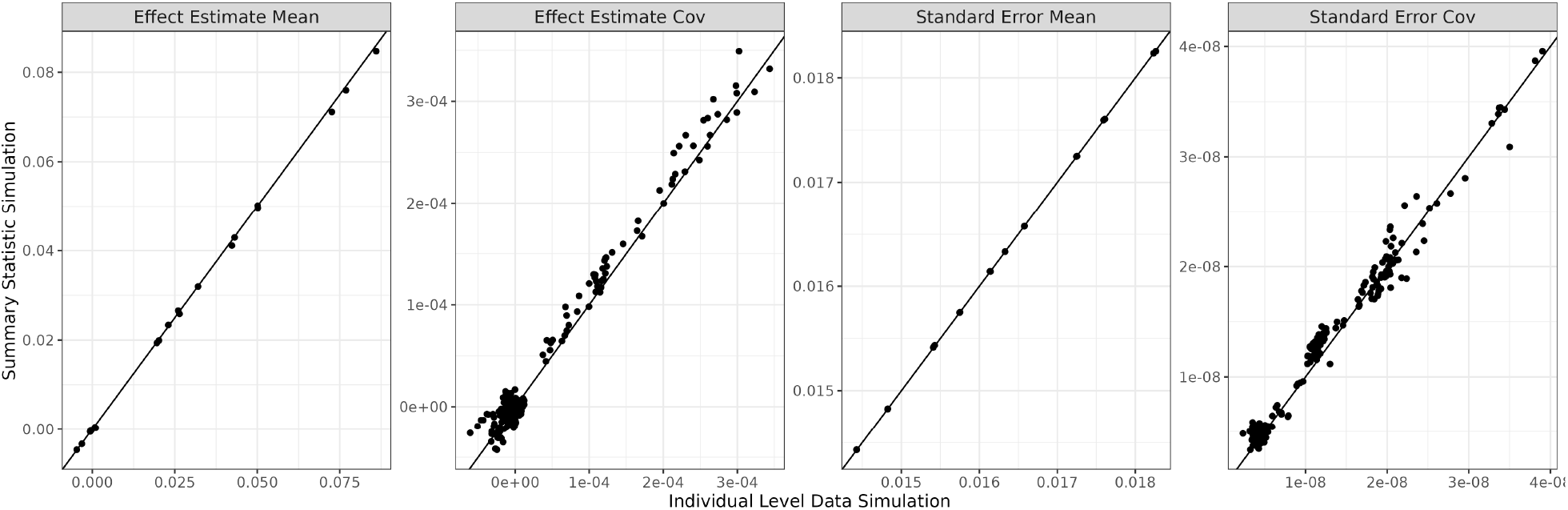
Comparison of summary statistics from the full data simulation method and the direct summary statistic simulation method used by `GWASBrewer`. Panels compare the mean and covariance of simulated effect standard error estimates. For mean plots, each point corresponds to one variant-trait pair. For covariance plots, each point corresponds to one pair of variant-trait pairs.

## 6 Examples

### 6.1 Simulating Multiple Traits from a DAG

In this example, we simulated data that could be used to evaluate a Mendelian randomization (MR) method in the presence of heritable confounding. We simulate GWAS data for an exposure, *X*, and outcome, *Y*, variable with a common heritable cause, *U*, shown in Figure 3. We then use package functions to identify exposure instruments that can be used with Mendelian randomization. In this case, our DAG includes three traits but we only need summary statistics for the exposure and the outcome. We can avoid simulating unneeded summary statistics for *U* by setting the sample size to zero.

**Figure 3.**
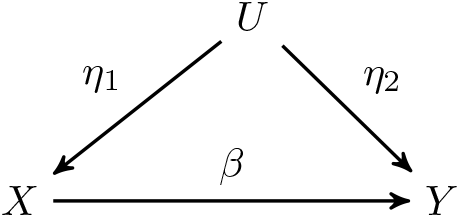
DAG for Mendelian randomization example.

We first set the direct effects matrix which is passed to the `G` parameter of `sim_mv`,

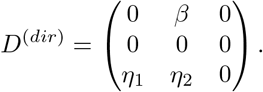

In this case, the traits are listed in the order of *X, Y, U*.

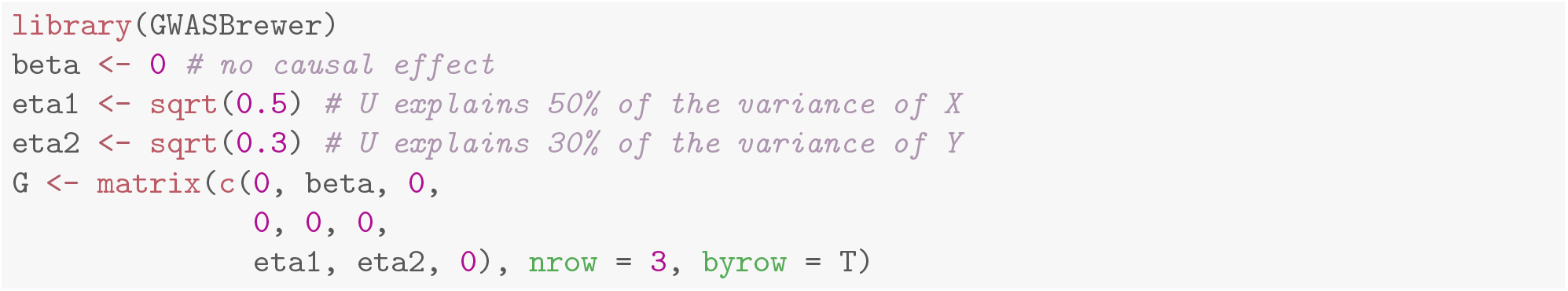

To make the simulations as realistic as possible, we generate data with LD and simulate the standard errors. We use the matrix format for sample size to indicate that there are 40,000 individuals in the exposure GWAS, 60,000 in the outcome GWAS, with 10,000 individuals overlapping. The last row and column of the sample size matrix are zero because we don’t need summary statistics for *U*.

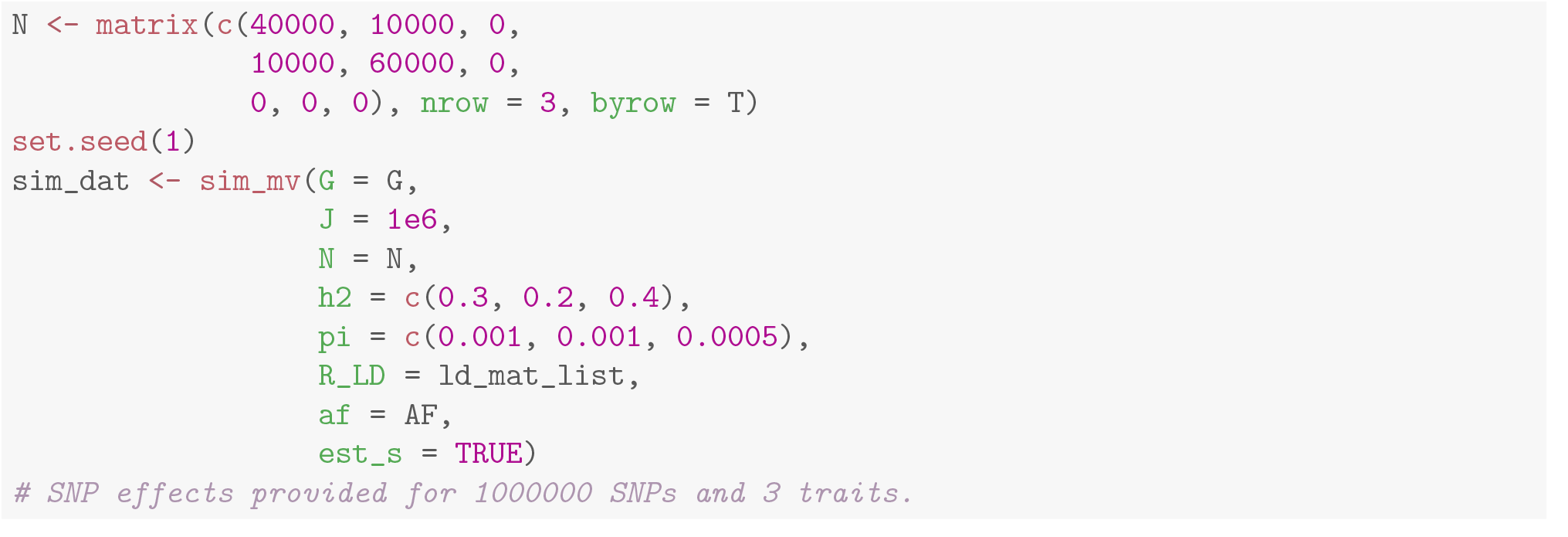

The summary statistics that we need are stored in `beta_hat` and `s_estimate`. No summary statistics are produced for *U*, because the sample size for *U* was given as 0.

In a typical MR analysis, we select the top independent variants associated with the exposure to use as instruments. We can do this using the `sim_ld_prune` function. Below, we use this function to LD prune using the *p*-value for *X*, the first trait (`pvalue = 1`). The function returns a vector of variant indices.

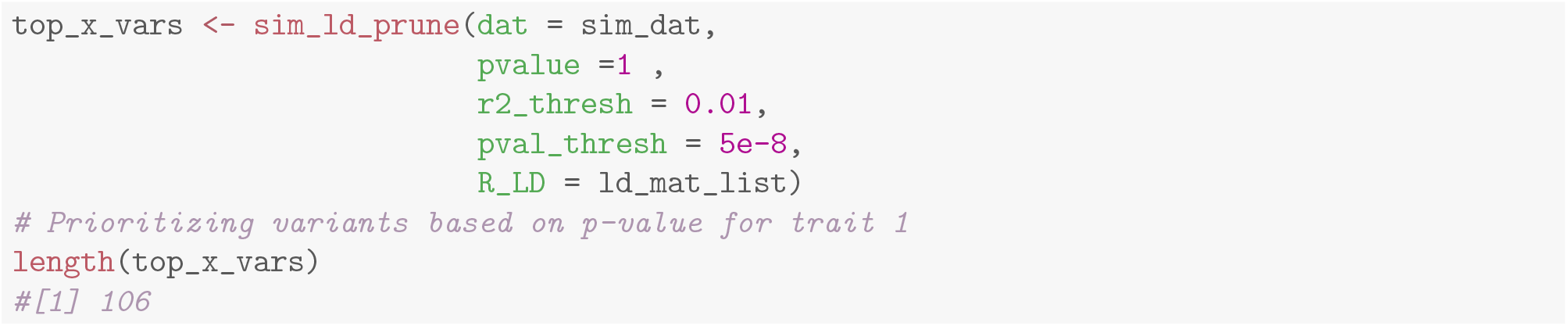

Alternatively, we could simulate a three sample MR study, in which there are two separate GWAS for *X*, one for selection and one for testing. To generate data for a third selection sample, we can use the `resample sumstats` function. For the selection GWAS, we only need data for *X* so we set sample sizes for the other two traits to zero. In this case we will use the same LD pattern for the selection GWAS as for the testing GWAS, but we could have chosen to do otherwise. Alternatively, we could generate all three GWAS in a single call to `sim_mv` by creating a fourth trait *X*^*′*^ and setting the effect of *X* on *X*^*′*^ to 1. This method would not allow the selection GWAS to have a different LD pattern from the other studies but would allow a partially overlapping cohort.

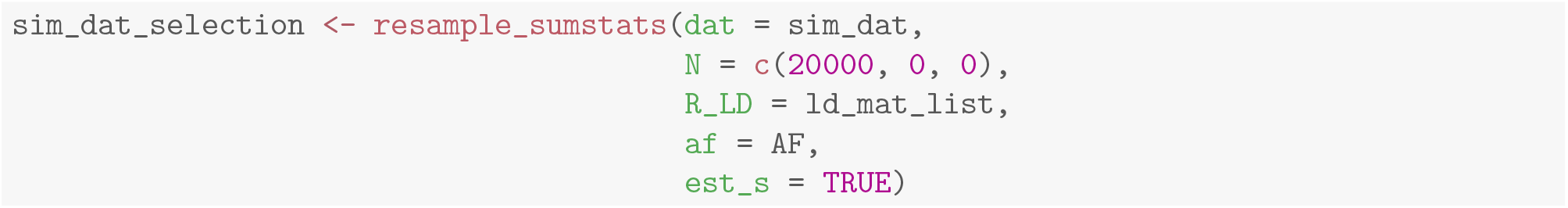

### 6.2 Advanced Effect Size Options

In this example, we demonstrate some advanced options for specifying the distribution of variant effects. These are applicable to scenarios which require more control over the location of causal variants or the distribution of causal variant effect size than provided by the default options. For example several alternative heritability models have been proposed including the LDAK[3] and LDSC [1] models. Under the LDSC model, expected SNP heritability is constant across variants, while in the LDAK model 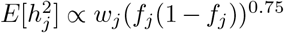, where 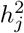 is the heritability contributed by variant *j*, and *w*_*j*_ is a weight that depends on the LD score of the variant. Proposals have also been made for estimating heritability enrichment by variant annotations [4, 2]. To evaluate these methods in simulations, we would need to simulate data with different effect distributions, potentially dependent on variant annotations.

#### 6.2.1 Controlling the Location of Causal Variants

The location of causal variants is controlled through the `pi` parameter which can be given as a scalar, a vector, or a matrix. Below, we show an example in which all causal variants are restricted to the first half of the genome. Here we use `G = 1`, to simulate one trait.

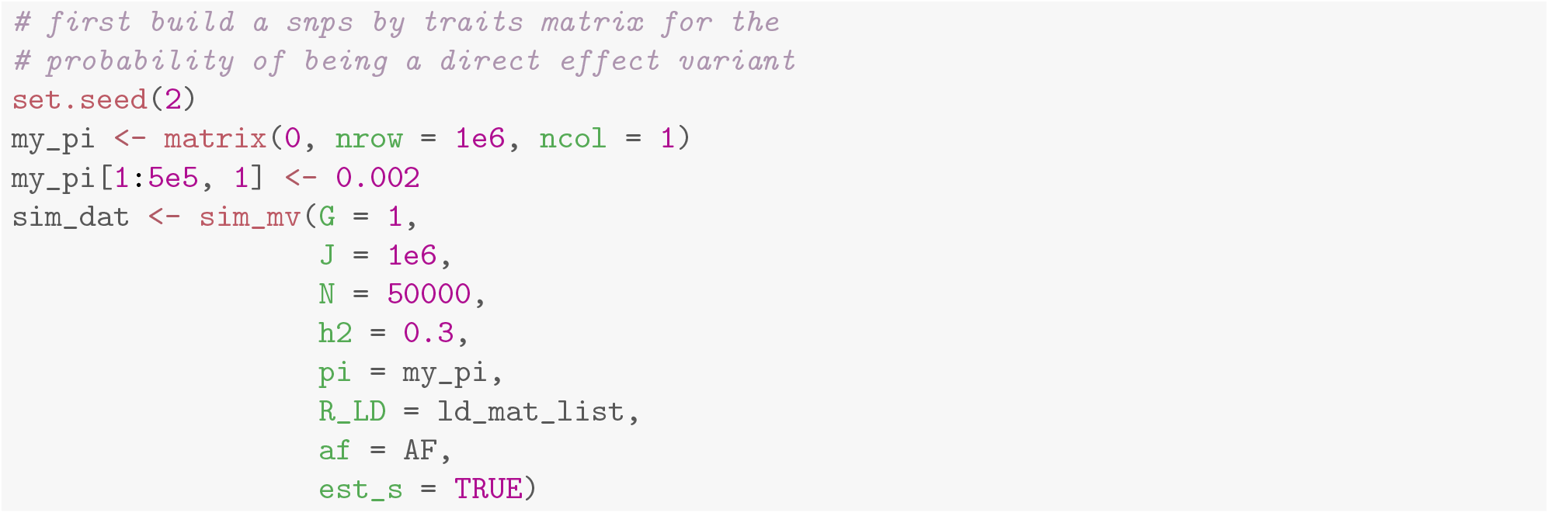

In a more complex example, we let the probability that a variant is causal be proportional to its LD score:

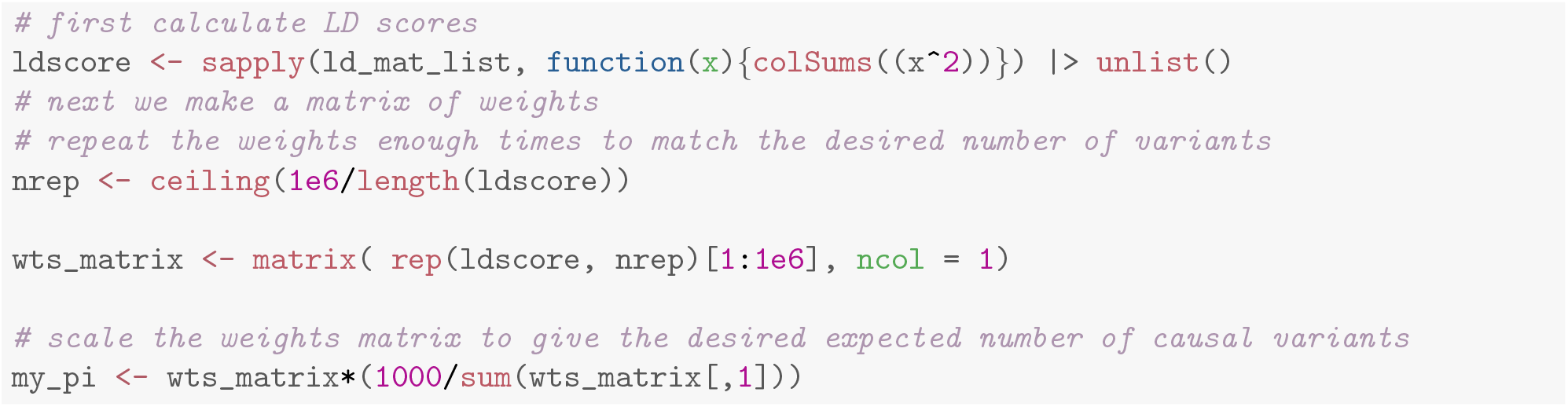

The weight matrix `my pi` would then be passed to the `pi` argument of `sim_mv`.

#### 6.2.2 Controlling the Distribution of Variant Effects

The distribution of direct effects can be controlled with the `snp_effect_function` argument. This argument can accept a single function that will be used for all traits or a list of *K* functions. Fucntions passed to `snp_effect_function` should accept three parameters: `n`, `sd`, and `snp_info` and return a vector of `n` standardized effect sizes (*a*_1_, …, *a*_*n*_) such that 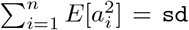. Many simple distribution functions will ignore the `snp_info` argument, which can be used to allow effect sizes to depend on an annotation or the allele frequency.

`GWASBrewer` contains two helper functions for forming variant effect functions for common distribution families, the scale mixture of normals (`mixnorm_to_scale_fam`), and effects specified directly up to a constant (`fixed_to_scale_fam`). In the scale mixture of normals, the direct effect of variant *j* on trait *k* is drawn from a distribution

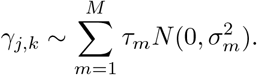

The set of mixture parameters *τ*_1_, …, *τ*_*M*_ and variance parameters 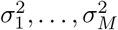 specify the distribution. To generate causal effects from a two-part mixture in which 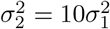 with 90% of causal variants drawn from the first component and 10% from the second, we use

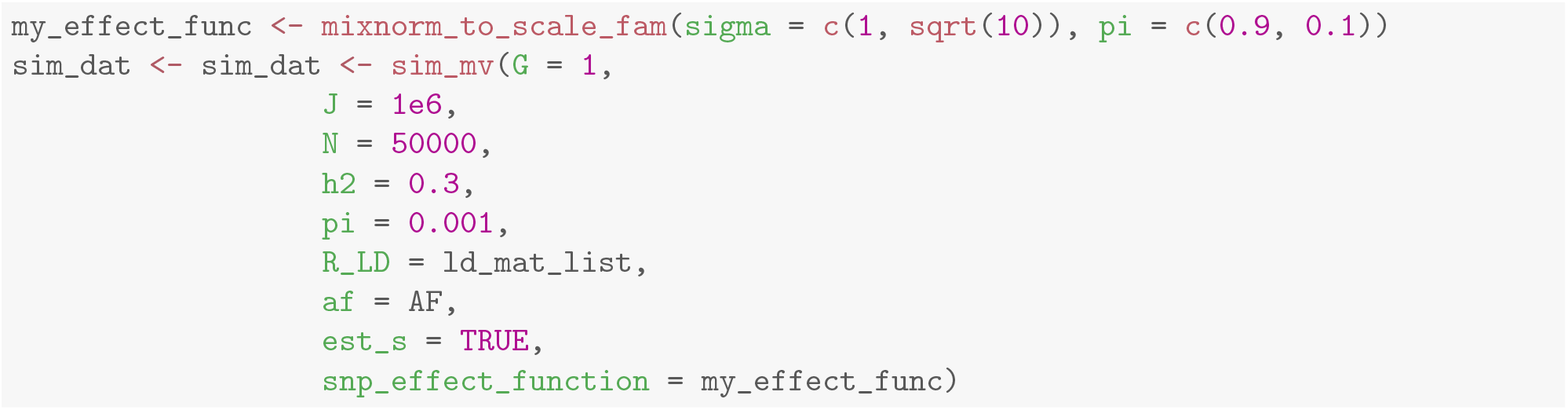

Note that there are generally multiple ways to get the same result. In example above, 0.1% of variants are causal. We could achieve equivalent results by including a 0 component in the scale mixture of normals and setting `pi` to 1.

Alternatively, the user can write their own variant effect functions. Here we show an example in which effect size depends on two annotations, mimicking a simplified version of the LDAK annotation mo del. We generate data in which

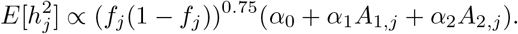

For our example, annotations *A*_1_ and *A*_2_ will be generated randomly. In a real application, these annotations could be real genomic annotations such as chromatin accessibility or distance to promoter. Annotations are supplied to `sim_mv` through the `snp_info` argument. These are passed to the variant effect function with an added column, `AF` containing allele frequency. We first write a variant effect function to give the desired distribution using *α*_0_ = 0.9, *α*_1_ = 0.1, and *α*_2_ = 0.13.

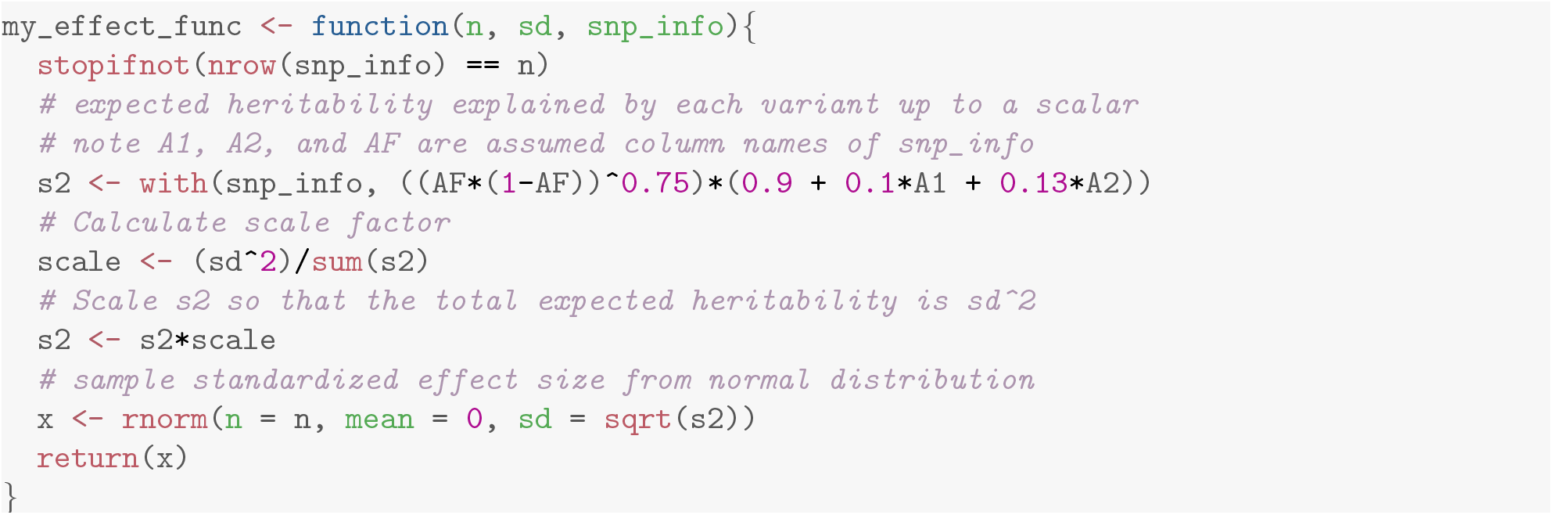

Next we generate random annotations. If we are using LD, the number of rows in the annotation data frame should match the total size of the LD pattern.

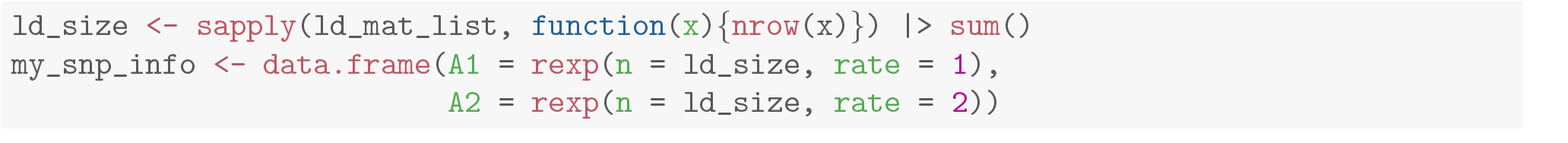

Finally, we can run `sim_mv` passing `my_effect_func` to `snp_effect_function` and `my_snp_info` to `snp_info`.

### 6.3 Simulating Individual Level Data

In the last example, we simulate data that could be used to evaluate accuracy of a polygenic risk score (PRS) constructed from GWAS results. This simulation demonstrates use of the `resample inddata` to generate individual level genotype and phenotype data.

First, we simulate summary statistics for the discovery GWAS that will be used to build the PRS.

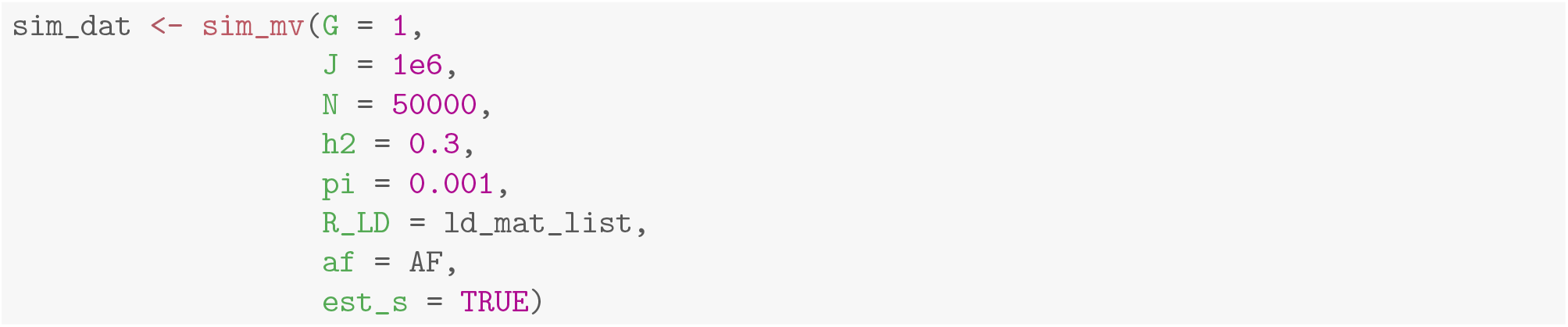

After constructing the PRS using any method, we need to simulate testing data. We will use the same LD structure used to generate discovery data. Using a different LD structure would allow us to measure cross-ancestry PRS transportability under the assumption that only LD and not effect sizes differ between populations. To generate indivdual level data, we use the `resample_inddata` function. The code below generates 1000 test samples. By default, `resample_inddata` uses `hapsim`[16] to simulate genotypes. However, alternative genotype simulation functions can be supplied to the `sim_func` argument.

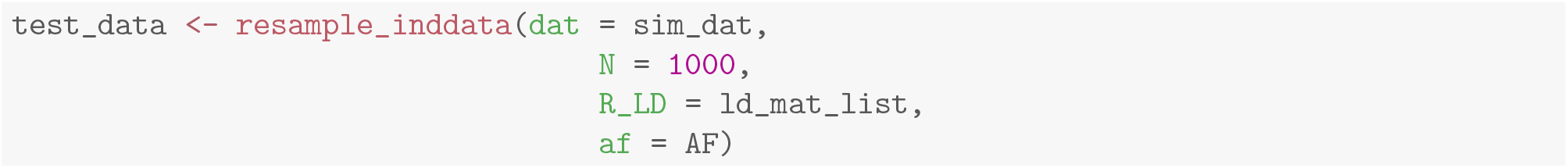

In a simulation study, we would need to repeat each of these steps many times. This will be more efficient if we generate only one set of test genotypes to be used in every iteration. To facilitate this, `resample_inddata` has two alternative modes, one in which it generates genotype data only, and one in which it accepts genotype data and simulates only phenotypes.

To generate only genotype data, we simply omit the `dat` argument and add the `J` argument to specify the desired number of variants to generate.

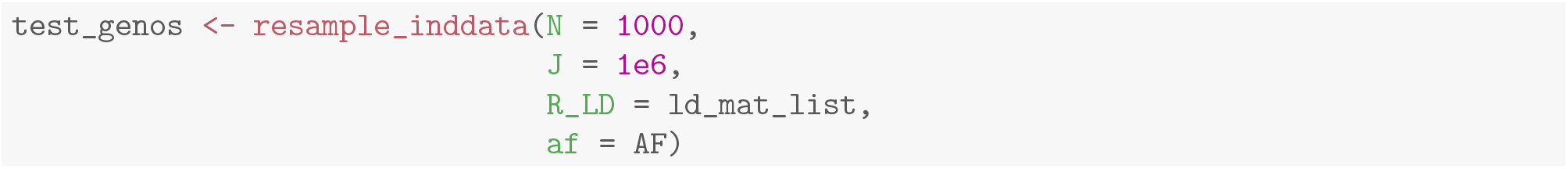

To generate phenotype data only, we can pass in the previously generated genotypes.

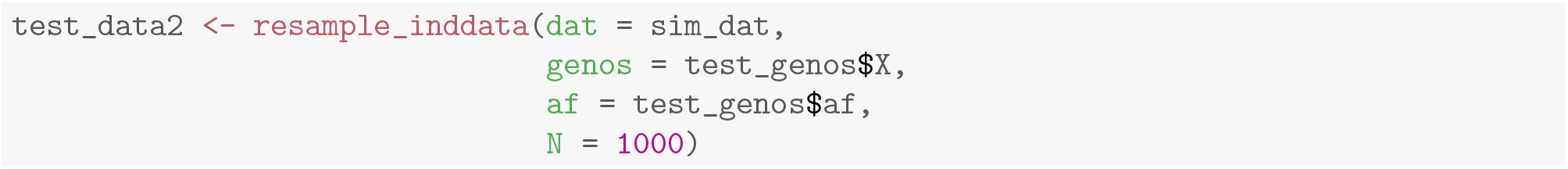

## 7 Discussion

We have introduced `GWASBrewer`, a software package for efficiently simulating GWAS summary data for multiple traits, allowing for LD between variants, correlation between traits, and any amount of sample overlap. We demonstrated that summary statistics produced by `GWASBrewer` have the same distribution as summary statistics produced through individual level data simulation. Simulating summary statistics directly can many times faster than simulating individual level genotype and phenotype data and requires less data storage. As a result, `GWASBrewer` can facilitate more in-depth and realistic simulations, leading to more accurate characterizations of method performance. `GWASBrewer` has potential utility for evaluating a wide range of statistical methods, including Mendelian randomization, colocalization, and heritability estimation methods. `GWASBrewer` allows for flexible specification of effect size distributions, and provides useful utility functions to aid in downstream evaluation of data. `GWASBrewer` also provides a unique feature of simulating standard error estimates, rather than returning only the true standard error of simulated effect sizes.

Although we have attempted to design `GWASBrewer` to be flexible enough to address many simulation needs, it has several limitations, some of which we hope to address in future releases. `GWASBrewer` currently only supports continuous traits related through linear structural equation models, and does not model gene-environment or gene-gene interactions. Additionally, `GWASBrewer` does not include confounding bias from factors such as uncorrected population structure, and generates data only for unrelated samples. In future releases, we plan expand support for binary traits, with linear relationships specified on the liability scale and expand the error model to allow for residual confounding.

## Supporting information

R script for simulations

## 8 Supplementary Note

### 8.1 Simulation of Standard Errors

In this section we describe the strategy used by `GWASBrewer` to simulate standard error estimates. We begin with two useful results.

#### Lemma 8.1.

*Let* **X**_1_, …, **X**_*N*_ *be an IID sample from a d-variate distribution with mean μ and at least four finite moments. Let* 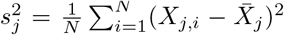 *be the sample variance in the jth dimension, let* 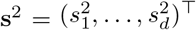, *and let* 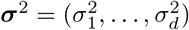 *be the d-vector of component-wise variances. Then*

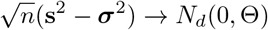

*where* Θ *is a d × d covariance matrix with elements*

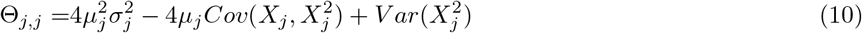

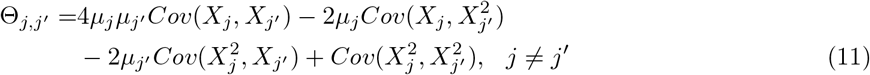

*Proof*. The proof of this lemma follows from the multivariate central limit theorem and the multivariate delta method. Without loss of generality, we derive the expressions above for *d* = 2. Let 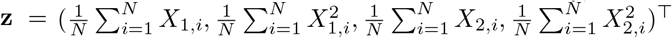 and let 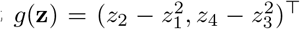 so that 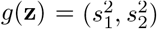. By the multivariate central limit theorem,

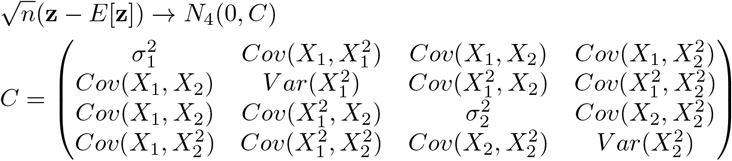

The gradient of the function *g* evaluated at *E*[**z**] is

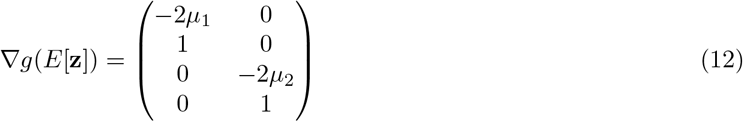

Thus, by the multivariate delta method,

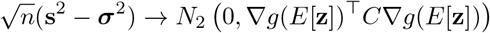

Some algebra shows that *∇g*(*E*[**z**])^*⊤*^*C∇g*(*E*[**z**]) is equal to Θ as specified above.

#### Corollary 8.1.1.

*Let X*_1_, …, *X*_*N*_ *be an IID sample from a one-dimensional distribution with at least four finite moments. Denote the mean, variance, and fourth central moment as μ, σ*^2^, *μ*_4_ *and let* 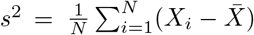. *Then*

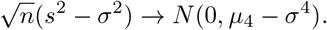

*Proof*. This is simply the one-dimensional version of the preceding lemma.

We start by describing simulation of standard error estimates for one trait. Let ŝ be the *J*-vector of standard error estimates for all variants. Using the expression in (2), ŝ_*j*_ can be written as

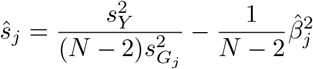

where 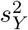and 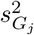 are the sample variances of the phenotype *Y* and variant *G*_*j*_ respectively. We make the simplifying assumption that 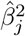 is independent of 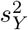 and 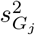 and approximate the distribution of *ŝ*_*j*_ as

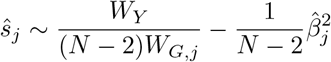

where *W*_*Y*_ and *W*_*G,j*_ are respectively samples from the asymptotic distributions of 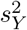 and 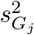. Specifically, to sample from the asymptotic distribution of 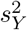, we apply the corollary above, assuming that the fourth moment of the phenotype distribution is equal to the fourth moment of a *N* (0, 1) distribution, or 3. Thus, we sample a single realization of *W*_*Y*_ *∼ N* (0, 2). This single draw is shared for all variants because estimates for all variants are made using the same set of individuals.

For the denominator, we need to sample from the distribution of, 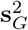, the vector of sample variances for all *J* variants. Here we apply the full multivariate version of the Lemma. We assume that the *J*-vector of genotypes for individual *i* is the sum of two independent vectors of Bernoulli random variables (a paternal and maternal haplotype), **G**_*i*_ = **W**_*i*,1_ + **W**_*i*,2_, where **W**_*i,k*_ (*k* = 1, 2) are IID *J*-dimensional vectors of correlated

Bernoulli random variables such that *W*_*i,k,j*_ *∼ Bernoulli*(*f*_*j*_) and *Cor*(*W*_*i,k,j*_, *W*_*i,k,j*_^*′*^) = *ρ*_*j,j*_^*′*^ as specified by the LD matrix, for *k ∈ {*1, 2*}*. From this distribution, we can calculate that

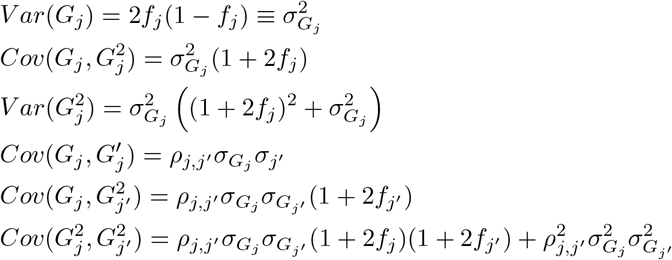

Plugging these in to the expressions in (10) and (11) and condensing terms, we obtain

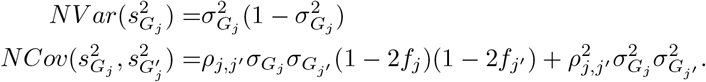

This allows us to sample **W**_*G*_ from the normal distribution defined by these expressions using user-specified allele frequencies and LD matrix.

The procedure for sampling standard errors for *K* traits is the same except that we now sample 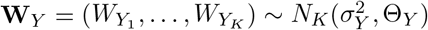, with one element for each trait. If there is sample overlap, then Θ_*Y*_ is not diagonal and we again apply the multivariate version of the Lemma 8.1. In this case, since all phenotypes have mean zero, we have

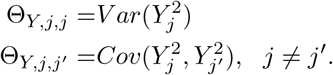

As previously, we use a normal distribution to select values for 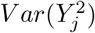 and 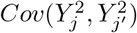. This gives 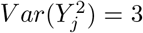 and 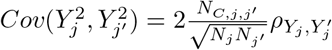 where *ρ*_*Yj*_, 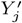 is the observational correlation of *Y*_*j*_ and 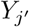 and *N*_*C,j,j*_^*′*^ is equal to the number of samples overlapping between the *j* and *j*^*′*^ GWAS study.

### 8.2 Additional Simulation Accuracy Results

Figures 4,5,6, and 7 show quantile-quantile plots comparing the distribution of effect estimates and standard error estimates and products of these to distributions obtained by simulating summary statistics directly. These plots all support they hypothesis that summary statistics generated by `GWASBrewer` have the same distribution as summary statistics obtained by first simulating individual level genotype and phenotype data. R code implementing these simulations is included in the Supplementary Materials.

**Figure 4.**
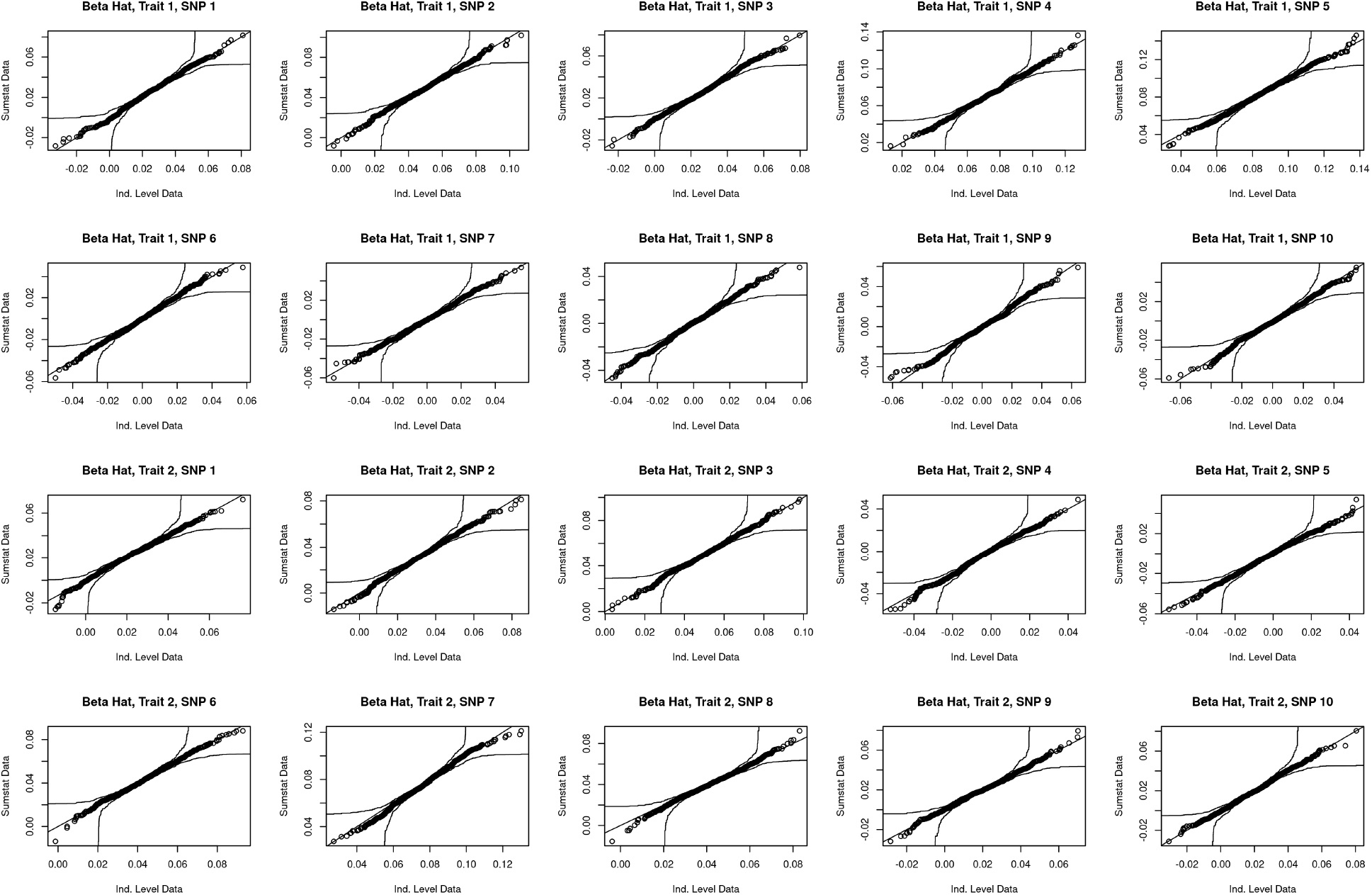
Q-Q plots comparing the distribution of effect estimates sampled directly and generated from individual-level data for each trait-variant combination.

**Figure 5.**
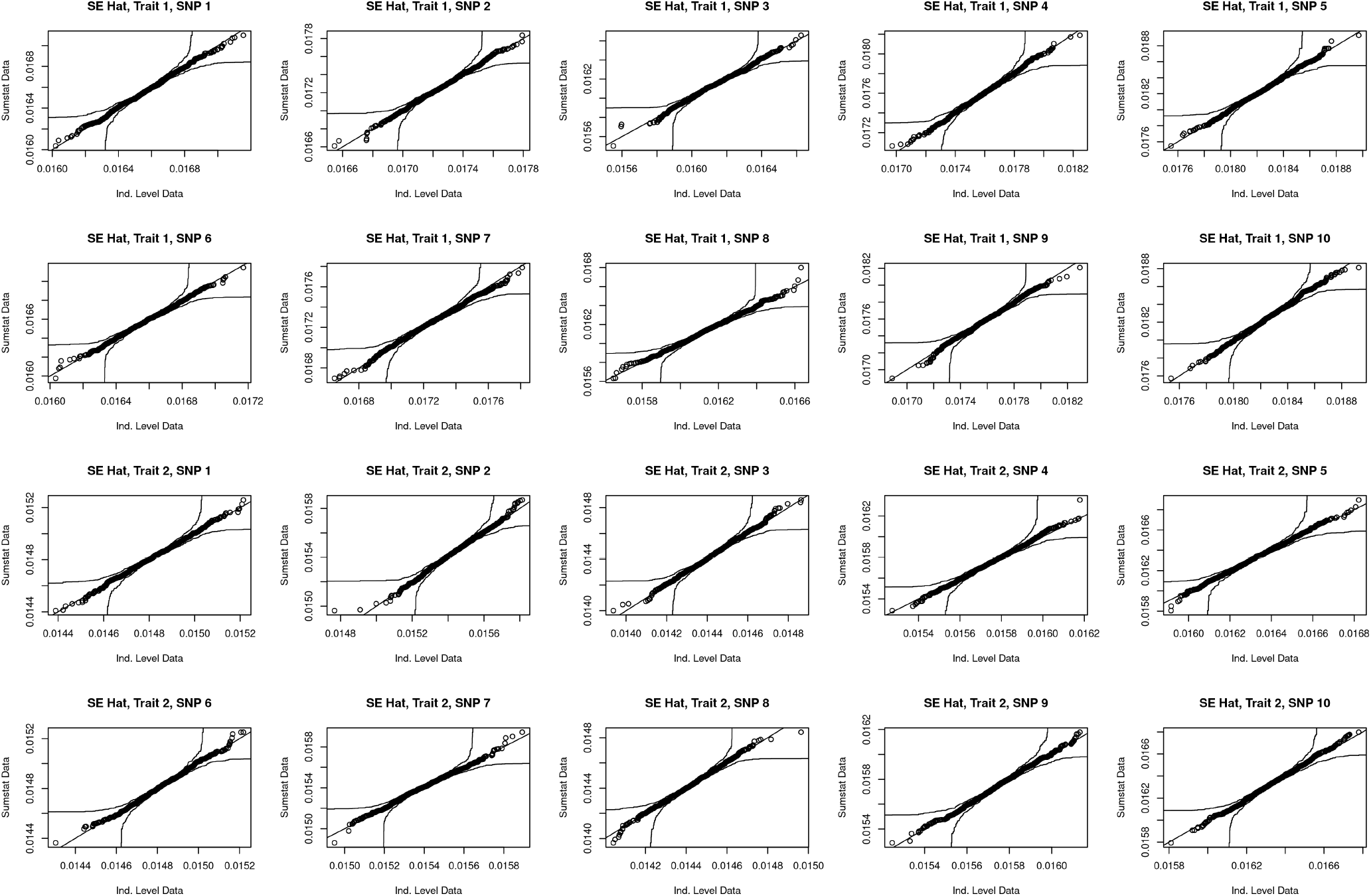
Q-Q plots comparing the distribution of standard estimates sampled directly and generated from individual-level data for each trait-variant combination.

**Figure 6.**
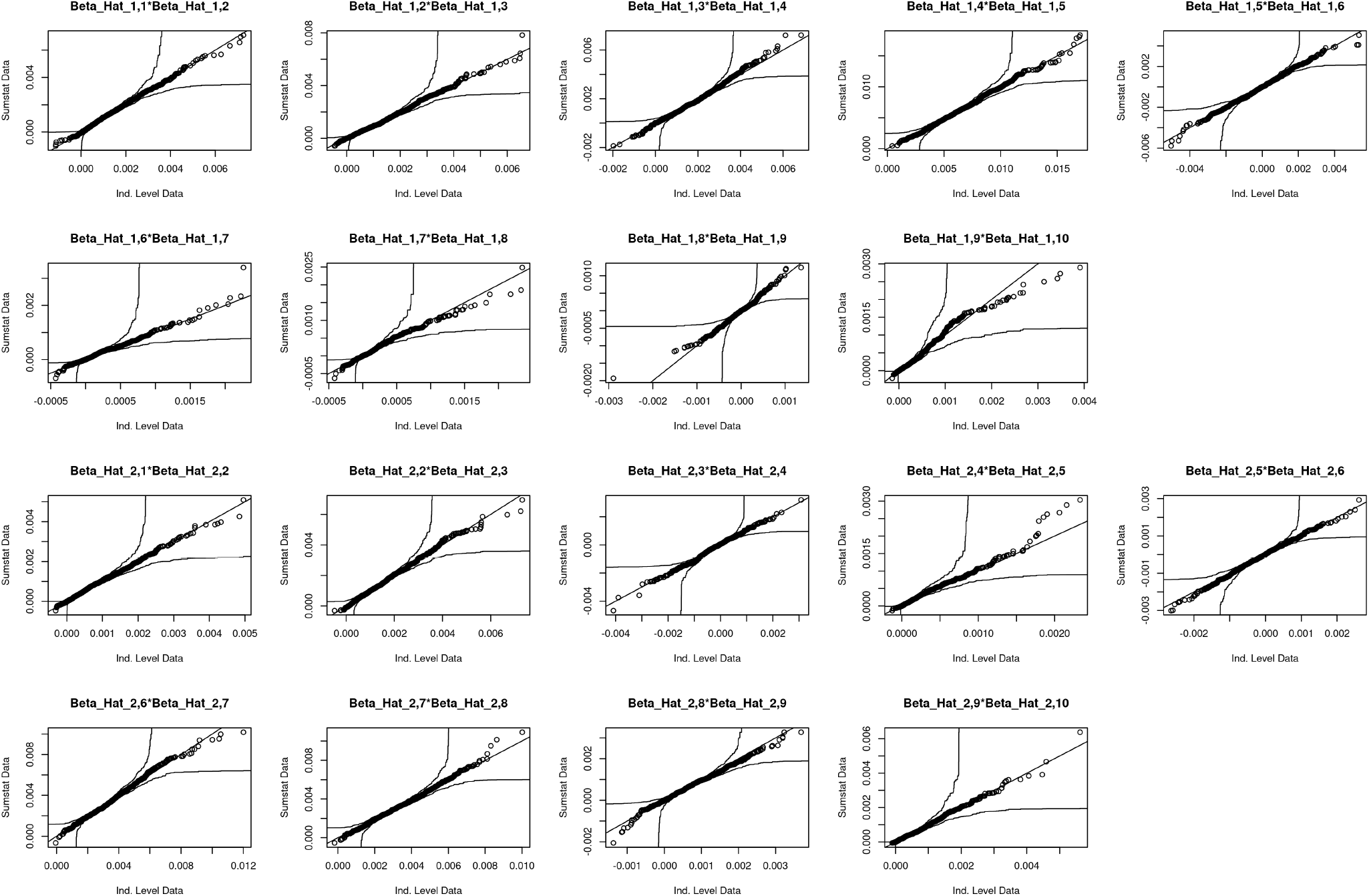
Q-Q plots comparing the distribution of products of effect estimates sampled directly and generated from individual-level data. Beta Hat i,j indicates the effect estimate for trait i and variant j.

**Figure 7.**
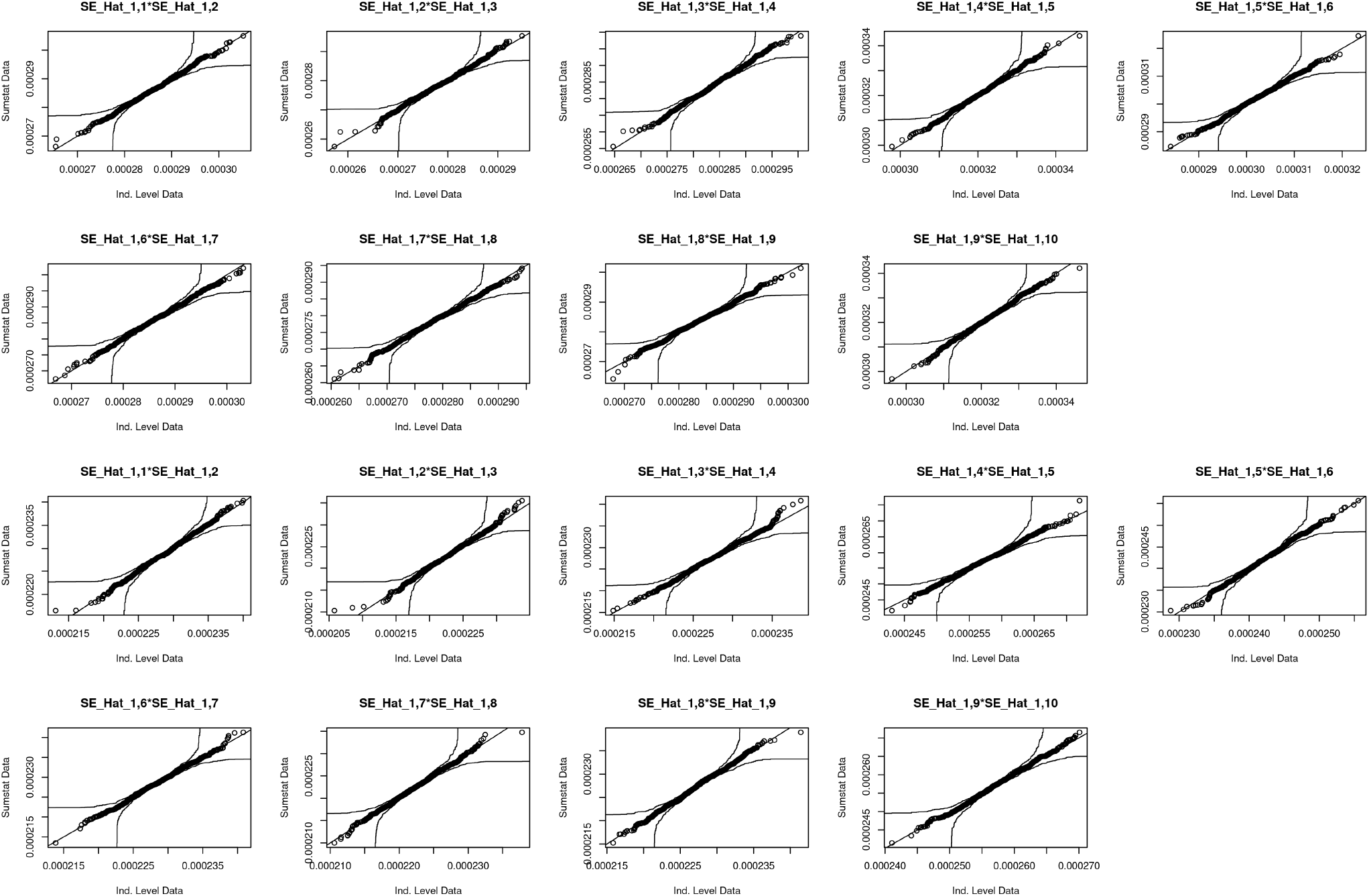
Q-Q plots comparing the distribution of products of standard error estimates sampled directly and generated from individual-level data. SE Hat i,j indicates the standard error estimate for trait i and variant j.

### 8.3 Inputs and Outputs of sim_mv

#### 8.3.1 Input Parameters

##### Arguments describing the traits

`G` : *K × K* matrix of direct trait effects (*D*^(*dir*)^).

`h2` : Expected trait heritability. This can be a scalar in which case all traits have the same heritability or a vector of length *K*.

`R obs` : Optional *K × K* matrix giving the observational correlation between traits. If missing, the correlation is calculated assuming independent direct environmental contributions.

`R_E` : Optional *K × K* matrix giving the correlation between environmental components of traits. Only one of `R obs` and `R_E` should be specified. Note that this is not the same as the correlation between the direct environmental components of traits.

##### Arguments describing GWAS study design

`N` : GWAS sample size. There are four accepted formats for `N`, scalar, vector, matrix, and dataframe. If `N` is a scalar, all GWAS have the same sample size and there is no sample overlap. If `N` is a vector, there is no sample overlap and each element gives the sample size for the corresponding trait. The matrix and dataframe formats can be used to specify sample overlap. If `N` is a matrix then `N[i,j]` specifies the number of samples in both study `i` and study `j` with `N[i,i]` giving the total sample size of study Finally, `N` can have dataframe format. In dataframe format, `N` should have columns named `trait 1` … `trait [K]` and `N`. The `trait [k]` columns will be interpreted as logicals and `N` gives the number of samples in each combination of studies (see examples below for more details). For `sim_mv`, matrix and dataframe format contain equivalent information. However, the dataframe format is useful with other package functions.

`J` : Total number of variants to simulate.

`R LD` : Optional list of LD blocks. `R LD` should be a list with each element describing a block in the block-diagonal LD matrix. Each element can be a matrix, a sparse matrix, or an eigen-decomposition. All elements should be correlation matrices, meaning that they have 1 on the diagonal and are positive definite.

`af` : Optional vector of allele frequencies. `af` can be a scalar, vector or function. If `af` is a function it should take a single argument, `n`, and return a vector of `n` allele frequencies. If `R LD` is supplied, `af` must be a vector with length equal to the size of the supplied LD pattern. Otherwise, `af` can be a scalar or vector of length *J*.

##### Arguments describing effect size distribution

`pi`: The probability that each variant has a direct effect on each trait. `pi` can be a scalar, vector of length *K*, or *J × K* matrix. If `pi` is a vector, each element corresponds to one trait. If `pi` is a matrix, each element corresponds to one variant-trait pair.

`sporadic pleiotropy`: Logical argument controlling whether or not variants can have direct effects on multiple traits. If `TRUE` (default) causal variants are chosen for each trait independently with no restrictions. If `FALSE`, causal variants are chosen for each trait in order and are not allowed to directly affect multiple traits. This option must be set to `TRUE` if `pi` is a matrix. In some cases, using `sporadic pleiotropy = FALSE` is inconsistent with other arguments, generating an error. For example if `pi` were set to 1, it would not be possible to avoid variants with causal effects on multiple traits.

`pi exact` Logical argument controlling whether there is variation in the number of direct effect variants. This argument defaults to `FALSE`, in which case the number of causal variants will be random. If set to `TRUE`, the number of direct effect variants for trait *k* will be exactly equal to `round(J * pi[k])`. If `pi` is a matrix, this argument must be `FALSE`

`snp effect function`: An optional user-specified function or list of functions to generate direct effect sizes. SNP effect functions should take arguments `n`, `sd`, and `snp info` and return a vector of `n` values with standard deviation `sd`. A function that does not return values with the specified variance (for example a deterministic function) can be used if `h2 exact = TRUE`. When this function is called by `sim_mv`, the `snp info` argument will be given a data frame including the allele frequency and any variant annotations supplied to the `snp info` argument of this function.

`snp info` Optional dataframe of variant information to be passed to variant effect functions. If `R LD` is specified, `snp info` should have number of rows equal to the size of the supplied LD pattern. Otherwise `snp info` should have `J` rows.

`h2 exact` : Logical argument controlling whether the realized heritability is exactly equal to the specified expected heritability.

##### Additional arguments

`est s` Logical argument. If `TRUE`, the function will sample estimates of standard errors. Defaults to `FALSE`.

#### 8.3.2 Output Object

`sim_mv` produces an object of class `sim_mv`. Below, we describe each of the elements of a `sim_mv` object.

##### Simulated summary statistics

`beta hat` : A *J × K* matrix of simulated effect estimates.

`s estimate` : If `est s` was set to true, this contains a *J × K* matrix of simulated standard errors

##### True variant effects

`beta marg` : True marginal variant associations, equal to the expected value `beta hat`.

`beta joint` : True joint variant effects. If `R LD` was missing then all variants are independent so `beta joint` and `beta marg` are equal.

s`e beta hat` : True standard errors of `beta hat`.

`direct SNP effects marg` : True marginal direct variant effects.

`direct SNP effects marg` : True joint direct variant effects. If there are no causal effects between traits or there is only one trait, then `direct SNP effects marg` is equal to `beta marg` and `direct SNP effects joint` is equal to `beta joint`.

##### Other useful information

`direct trait effects` : *K × K* direct trait effects matrix, equal to `G` provided to `sim_mv` (this is *D*^(*dir*)^).

`total trait effects` : Total trait effects matrix, equal to *D*^(*tot*)^.

`trait cor` : Observational trait correlation.

`R` : Correlation of columns of `beta hat` (equal to the identity if there was no sample overlap).

`Sigma G` : *K × K* genetic variance-covariance matrix. Diagonal elements of `Sigma G` give the realized trait heritability.

`Sigma E` : *K × K* environmental variance-covariance matrix.

`snp info` : Dataframe of variant information. At a minimum, `snp info` contains allele frequencies of each variant. If `snp info` was provided to `sim_mv`, this dataframe also contains any annotations included there.

