## Supplementary material for "GWASBrewer: An R Package for Simulating Realistic GWAS Summary Statistics": R script for simulations: GWASBrewer_dist_sims.R

```
library(GWASBrewer)
library(hapsim)
run_test <- function(nrep, N_df, J, h2, R_E, R_LD = NULL, af, G,
                     mypi, effect_func){

  dat <- sim_mv(N = 0,
                J = J,
                h2 = h2,
                pi = mypi,
                G = G,
                R_LD = R_LD,
                R_E  = R_E,
                af = af,
                est_s = TRUE,
                snp_effect_function = effect_func, 
                h2_exact = TRUE)
  res <- replicate(n = nrep, expr = {
    ss_dat <- resample_sumstats(dat = dat,
                              N = N_df,
                              R_LD = R_LD,
                              af = af,
                              est_s = TRUE)
    gw_dat <- resample_inddata(dat = dat,
                              N = N_df,
                              R_LD = R_LD,
                              af = af,
                              calc_sumstats = TRUE)
    cbind(ss_dat$beta_hat, gw_dat$beta_hat, ss_dat$s_estimate, gw_dat$s_estimate)
  })
  return(list(res = res, dat = dat))
}

# 10 SNPs, 2 trait, sample overlap, with LD
N_df <- data.frame("trait_1" = c(1, 1, 0), "trait_2" = c(1, 0, 1),
                   N = c(6000, 2000, 4000))
my_pi <- matrix(0, nrow = 10, ncol = 2)
ix1 <- c(2, 5)
my_pi[ix1, 1] <- 1
ix2 <- c(3, 7,9)
my_pi[ix2, 2] <- 1
R_E <- matrix(c(1, 0.7, 0.7, 1), nrow = 2)
A <- readRDS("sandbox/A.RDS")
# A
# [,1]  [,2]  [,3] [,4]  [,5]
# [1,]  1.00  0.60  0.20 -0.1 -0.07
# [2,]  0.60  1.00  0.60 -0.1 -0.07
# [3,]  0.20  0.60  1.00 -0.1 -0.07
# [4,] -0.10 -0.10 -0.10  1.0  0.90
# [5,] -0.07 -0.07 -0.07  0.9  1.00
af <- c(0.35, 0.3, 0.4, 0.72, 0.75)
R_LD = list(A)
my_effect_func <- function(n, sd, ...){
  x <- c(1, 2, 3)
  x <- x[(seq(n) %% 3) + 1]
}

set.seed(113)
res5 <- run_test(1000, N_df, J= 10, h2 = 0.004, R_E = R_E , R_LD = R_LD,
                 af = af, G = matrix(0, nrow = 2, ncol =2),
                 mypi = my_pi, effect_func = my_effect_func)
#saveRDS(res5, file = "~/Dropbox/Apps/Overleaf/simGWAS/2024-03-28_simple_dist_113.RDS")

res5 <- res5$res

beta_ss_means <- rowMeans(rbind(res5[,1,], res5[,2,]))
beta_gw_means <- rowMeans(rbind(res5[,3,], res5[,4,]))
res_sum <- data.frame(full_data = beta_gw_means,
                  ss_data = beta_ss_means,
                  type = "Effect Estimate Mean")
#plot(beta_ss_means, beta_gw_means)
#abline(0, 1)

beta_ss_sd <- apply(rbind(res5[,1,], res5[,2,]), 1, sd)
beta_gw_sd <- apply(rbind(res5[,3,], res5[,4,]), 1, sd)
res_sum <- bind_rows(res_sum,
                     data.frame(full_data = beta_gw_sd,
                                ss_data = beta_ss_sd,
                                type = "Effect Estimate SD"))

#plot(beta_ss_sd, beta_gw_sd)
#abline(0, 1)

# beta_ss_cor <- cor(t(rbind(res5[,1,], res5[,2,])))
# beta_gw_cor <- cor(t(rbind(res5[,3,], res5[,4,])))
# res_sum <- bind_rows(res_sum,
#                      data.frame(full_data = beta_gw_cor[upper.tri(beta_gw_cor)],
#                                 ss_data = beta_ss_cor[upper.tri(beta_ss_cor)],
#                                 type = "Effect Estimate Cor"))
beta_ss_cov <- cov(t(rbind(res5[,1,], res5[,2,])))
beta_gw_cov <- cov(t(rbind(res5[,3,], res5[,4,])))
res_sum <- bind_rows(res_sum,
                     data.frame(full_data = beta_gw_cov[upper.tri(beta_gw_cov, diag =T)],
                                ss_data = beta_ss_cov[upper.tri(beta_ss_cov, diag = T)],
                                type = "Effect Estimate Cov"))
#plot(beta_ss_cor, beta_gw_cor)
#abline(0, 1)
#

s_ss_means <- rowMeans(rbind(res5[,5,], res5[,6,]))
s_gw_means <- rowMeans(rbind(res5[,7,], res5[,8,]))
res_sum <- bind_rows(res_sum,
                     data.frame(full_data = s_gw_means,
                                ss_data = s_ss_means,
                                type = "Standard Error Mean"))
#plot(s_ss_means, s_gw_means)
#abline(0, 1)

s_ss_sd <- apply(rbind(res5[,5,], res5[,6,]), 1, sd)
s_gw_sd <- apply(rbind(res5[,7,], res5[,8,]), 1, sd)
#plot(s_ss_sd, s_gw_sd)
res_sum <- bind_rows(res_sum,
                     data.frame(full_data = s_gw_sd,
                                ss_data = s_ss_sd,
                                type = "Standard Error SD"))
#abline(0, 1)

# s_ss_cor <- cor(t(rbind(res5[,5,], res5[,6,])))
# s_gw_cor <- cor(t(rbind(res5[,7,], res5[,8,])))
# res_sum <- bind_rows(res_sum,
#                      data.frame(full_data = s_gw_cor[upper.tri(s_gw_cor)],
#                                 ss_data = s_ss_cor[upper.tri(s_ss_cor)],
#                                 type = "Standard Error Cor"))
s_ss_cov <- cov(t(rbind(res5[,5,], res5[,6,])))
s_gw_cov <- cov(t(rbind(res5[,7,], res5[,8,])))
res_sum <- bind_rows(res_sum,
                     data.frame(full_data = s_gw_cov[upper.tri(s_gw_cov, diag =T)],
                                ss_data = s_ss_cov[upper.tri(s_ss_cov, diag = T)],
                                type = "Standard Error Cov"))
#plot(s_ss_cor, s_gw_cor)
#abline(0, 1)
res_sum$type <- factor(res_sum$type, levels = c("Effect Estimate Mean",
                                        "Effect Estimate SD",
                                        "Effect Estimate Cov",
                                        "Standard Error Mean",
                                        "Standard Error SD",
                                        "Standard Error Cov"))
s <- 12
p <-  res_sum %>%
      filter(! type %in% c("Effect Estimate SD", "Standard Error SD")) %>%
      ggplot() +
      geom_point(aes(x = full_data,
                     y = ss_data)) +
      geom_abline(slope = 1, intercept = 0) +
      xlab("Individual Level Data Simulation") +
      ylab("Summary Statistic Simulation") +
      theme_bw() +
      theme(axis.title = element_text(size = s),
            axis.text = element_text(size = s-2),
            strip.text = element_text(size = s)) +
      facet_wrap(~type, ncol = 4, scales= "free")

ggsave(p, file = "~/Dropbox/Apps/Overleaf/simGWAS/sim_results_main.png", height = 5, width = 15, units = "in", dpi = 300)


png("~/Dropbox/Apps/Overleaf/simGWAS/qq1.png", height = 10, width = 15, units = "in", res = 300)
par(mfrow = c(4, 5))
for(i in 1:10){
  qqplot(res5[i,3,], res5[i,1,], conf.level = 0.95, xlab = "Ind. Level Data", ylab = "Sumstat Data", main = paste0("Beta Hat, Trait 1, SNP ", i))
  abline(0, 1)
}

for(i in 1:10){
  qqplot(res5[i,4,], res5[i,2,], conf.level = 0.95, xlab = "Ind. Level Data", ylab = "Sumstat Data", main = paste0("Beta Hat, Trait 2, SNP ", i))
  abline(0, 1)
}
dev.off()

png("~/Dropbox/Apps/Overleaf/simGWAS/qq2.png", height = 10, width = 15, units = "in", res = 300)
par(mfrow = c(4, 5))
for(i in 1:10){
  qqplot(res5[i,7,], res5[i,5,], conf.level = 0.95, xlab = "Ind. Level Data", ylab = "Sumstat Data", main = paste0("SE Hat, Trait 1, SNP ", i))
  abline(0, 1)
}
for(i in 1:10){
  qqplot(res5[i,8,], res5[i,6,], conf.level = 0.95, xlab = "Ind. Level Data", ylab = "Sumstat Data", main = paste0("SE Hat, Trait 2, SNP ", i))
  abline(0, 1)
}
dev.off()

png("~/Dropbox/Apps/Overleaf/simGWAS/qq3.png", height = 10, width = 15, units = "in", res = 300)
par(mfrow = c(4, 5))
for(i in 1:9){
  qqplot(res5[i,3,]*res5[i + 1, 3,], res5[i,1,]*res5[i+1,1,], conf.level  = 0.95, 
         xlab = "Ind. Level Data", ylab = "Sumstat Data", main = paste0("Beta_Hat_1,", i, "*Beta_Hat_1,", i + 1))
  abline(0, 1)
}
plot.new()
for(i in 1:9){
  qqplot(res5[i,4,]*res5[i + 1, 4,], res5[i,2,]*res5[i+1,2,], conf.level  = 0.95, 
         xlab = "Ind. Level Data", ylab = "Sumstat Data", main = paste0("Beta_Hat_2,", i, "*Beta_Hat_2,", i + 1))
  abline(0, 1)
}
dev.off()

png("~/Dropbox/Apps/Overleaf/simGWAS/qq4.png", height = 10, width = 15, units = "in", res = 300)
par(mfrow = c(4, 5))
for(i in 1:9){
  qqplot(res5[i,7,]*res5[i+1,7,], res5[i,5,]*res5[i +1,5,], conf.level =  0.95, 
         xlab = "Ind. Level Data", ylab = "Sumstat Data", main = paste0("SE_Hat_1,", i, "*SE_Hat_1,", i + 1))
  abline(0, 1)
}
plot.new()
for(i in 1:9){
  qqplot(res5[i,8,]*res5[i+1,8,], res5[i,6,]*res5[i +1,6,], conf.level =  0.95, 
         xlab = "Ind. Level Data", ylab = "Sumstat Data", main = paste0("SE_Hat_1,", i, "*SE_Hat_1,", i + 1))
  abline(0, 1)
}
dev.off()
```
